# Endothelial restoration of CAD GWAS gene PLPP3 by nanomedicine suppresses YAP/TAZ activity and reduces atherosclerosis *in vivo*

**DOI:** 10.1101/2021.05.06.443006

**Authors:** Jiayu Zhu, Chih-Fan Yeh, Ru-Ting Huang, Tzu-Han Lee, Tzu-Pin Shentu, David Wu, Kai-Chien Yang, Yun Fang

**Author notes:** Equal contribution (Jiayu Zhu, Chih-Fan Yeh). Corresponding author: Yun Fang, **Email:**. **Author Contributions** J.Z., C.F.Y., R.T.H., T.H.L., T.P.S., and D.W. planned and executed experiments, analyzed data and interpreted results. K.C.Y. and Y.F. planned experiments and interpreted results. J.Z., C.F.Y., R.T.H., T.P.S., D.W, K.C.Y., and Y.F wrote and edited the manuscript. **Competing Interest Statement:**, There authors declare no competing interests.

## Abstract

Genome-wide association studies (GWAS) have suggested new molecular mechanisms in vascular cells driving atherosclerotic diseases such as coronary artery disease (CAD) and ischemic stroke (IS). Nevertheless, a major challenge to develop new therapeutic approaches is to spatiotemporally manipulate these GWAS-identified genes in specific vascular tissues *in vivo*. YAP (Yes-associated protein) and TAZ (transcriptional coactivator with PDZ-binding motif) have merged as critical transcriptional regulators in cells responding to biomechanical stimuli, such as in athero-susceptible endothelial cells activated by disturbed flow (DF). The molecular mechanisms by which DF activates while unidirectional flow (UF) inactivates YAP/TAZ remain incompletely understood. Recent studies demonstrated that DF and genetic predisposition (risk allele) of CAD/IS locus 1p32.2 converge to reduce phospholipid phosphatase 3 (*PLPP3*) expression in vascular endothelium. Restoration of endothelial *PLPP3 in vivo*, although remains challenging and unexplored, is hypothesized to reduce atherosclerosis. We devised a nanomedicine system integrating nanoparticles and *Cdh5* promoter-driven plasmids to successfully restore *PLPP3* expression in activated endothelium, resulting in suppressed YAP/TAZ activity and reduced DF-induced atherosclerosis in mice. Mechanistically, our studies discovered a molecular paradigm by which CAD/IS GWAS gene *PLPP3* inactivates YAP/TAZ by reducing lysophosphatidic acid (LPA)-induced myosin II and ROCK in endothelium under UF. These results highlight a new mechanistic link between GWAS and YAP/TAZ mechano-regulation and moreover, establish a proof of concept of vascular wall-based therapies employing targeted nanomedicine to manipulate CAD/IS GWAS genes *in vivo*.

## Main Text

Genome-wide association studies (GWAS) and subsequent mechanistic investigations have identified novel cell-specific molecular mechanisms causing atherosclerosis. A major challenge for GWAS-informed, new anti-atherosclerotic therapeutics is to spatiotemporally manipulate GWAS genes in specific tissues of interest *in vivo*. GWAS identified chromosome 1p32.2 as one of the most strongly associated loci with atherosclerotic diseases such as coronary artery disease (CAD) and ischemic stroke (IS). Our recent results (1, 2) highlighted phospholipid phosphatase 3 (*PLPP3*), located in 1p32.2, in endothelial mechano-transduction mechanisms associated with the focal nature of atherosclerosis in arterial curvatures and branches. While athero-protective unidirectional flow (UF) increases endothelial *PLPP3*, disturbed flow (DF) and CAD/IS risk allele at 1p32.2 converge to suppress *PLPP3* in endothelium prone to atherosclerosis. Endothelial *PLPP3* restoration *in vivo*, although remains challenging, is expected to lessen DF-induced atherosclerosis.

Yorkie homologues YAP (Yes-associated protein) and TAZ (transcriptional coactivator with PDZ-binding motif) recently emerged as critical transcriptional regulators in cellular mechano-transduction such as DF-induced endothelial activation at atherosclerosis-prone sites (3, 4). Regulatory mechanisms linking human genetics to flow-dependent YAP/TAZ activity have not been proposed. We first discovered that the CAD/IS GWAS gene *PLPP3* is a novel negative regulator of YAP/TAZ. *PLPP3* inactivates YAP/TAZ in vascular endothelium under UF. Suppression of UF-induced *PLPP3* using siRNA significantly increased YAP/TAZ activity, demonstrated by increased mRNA of YAP/TAZ downstream targets *CTGF* and *CYR61* (Fig. 1A) and increased YAP dephosphorylation and CYR61 protein (Fig. 1B). We hypothesized that YAP/TAZ inactivation by *PLPP3* is due to its phosphatase activity to hydrolyze lysophosphatidic acid (LPA). Indeed, treatments with Brp-LPA, a pan-LPA receptor antagonist and an inhibitor of Autotaxin, significantly abrogated the YAP dephosphorylation in *PLPP3* knock-downed HAEC (Fig. 1C). Inhibition of myosin II and ROCK by Blebbistatin and Y27632 respectively, increased phospho-YAP in *PLPP3* knock-downed HAEC (Fig. 1C). These data demonstrated for the first time, UF-induced *PLPP3* inactivates endothelial YAP/TAZ by reducing LPA-induced signaling and myosin II and ROCK activities.

**Figure 1.**
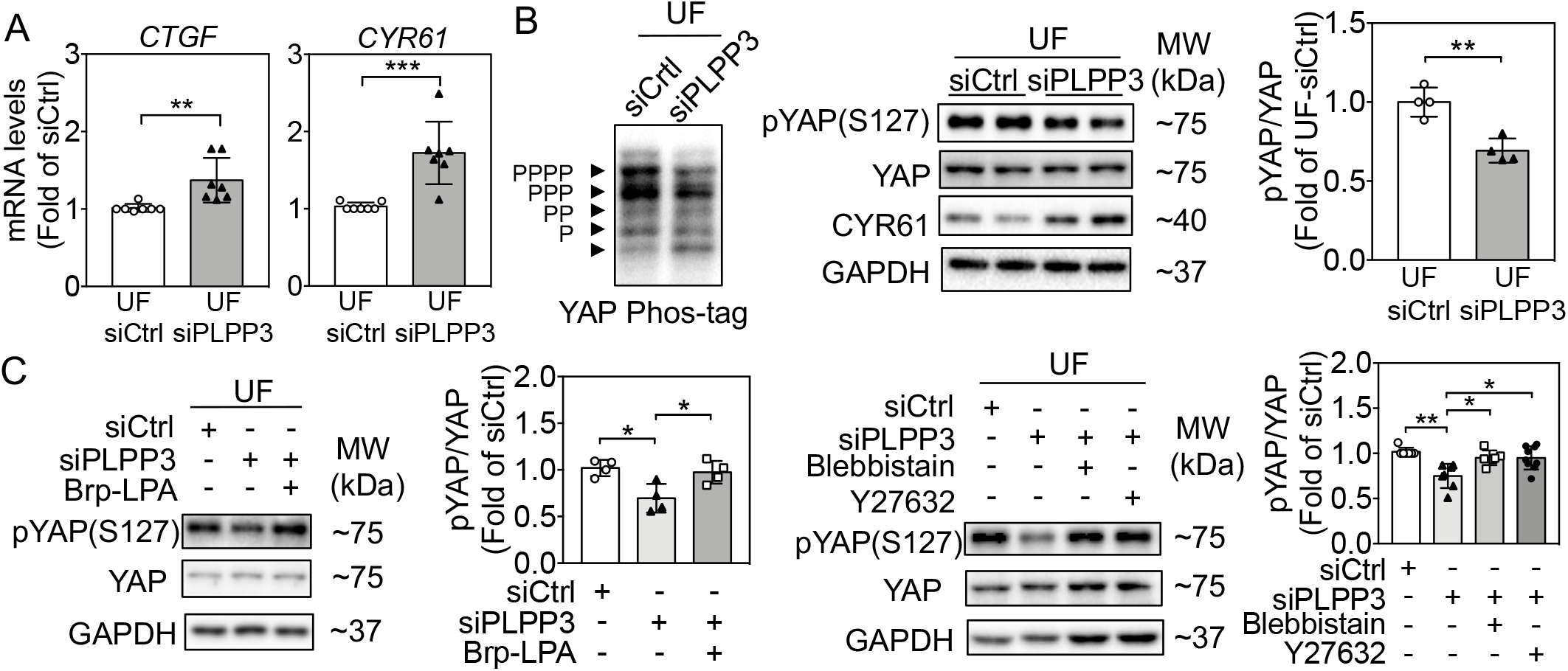
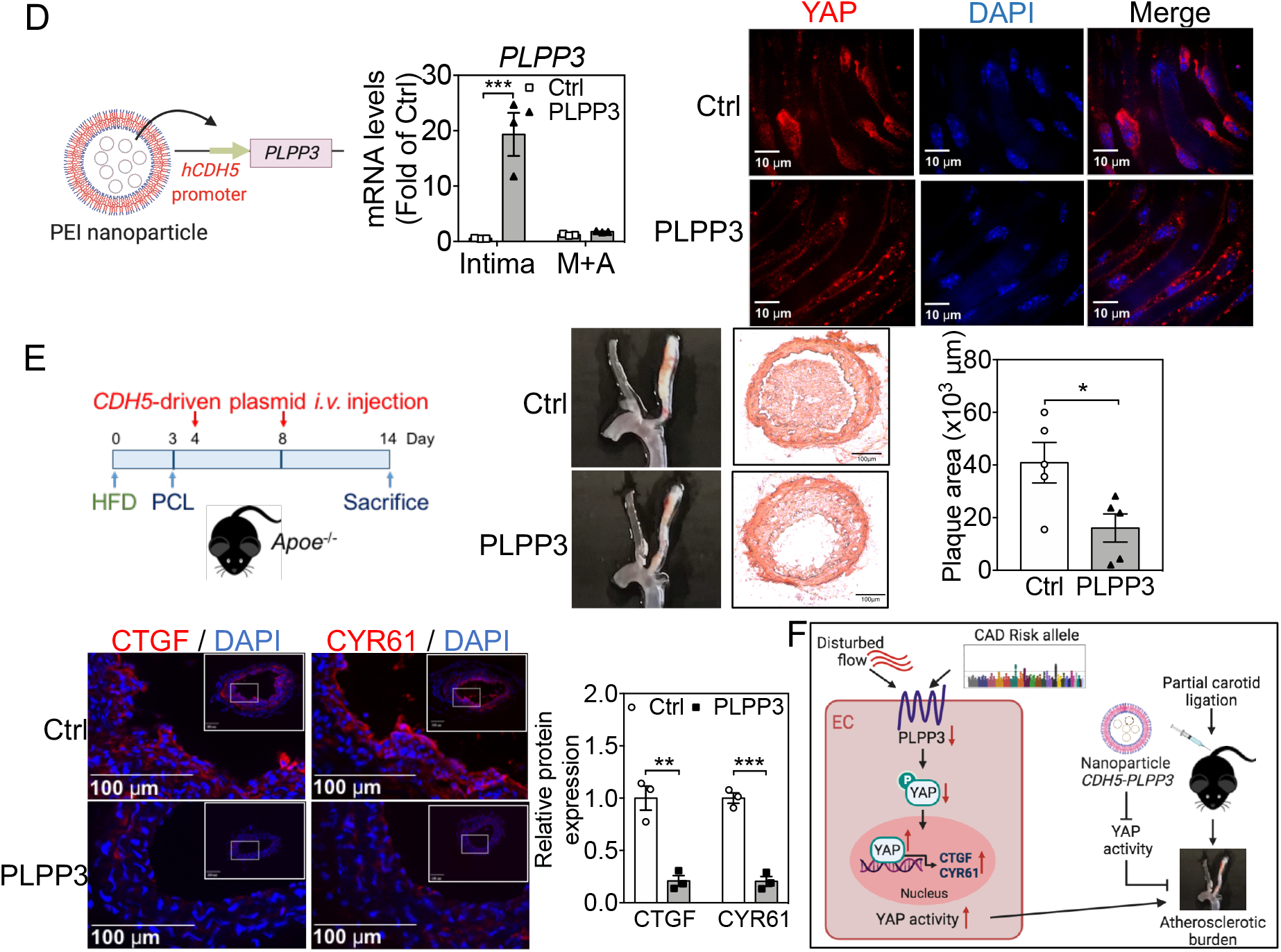
**(A)** Increased *CTGF* and *CYR61* mRNA by *PLPP3* knockdown in HAEC under UF (N=7). **(B)** Decreased YAP phosphorylation, reduced pYAP/YAP ratio, and induced CYR61 protein by *PLPP3* knockdown in HAEC under UF (N=4). **(C)** Increased pYAP/YAP ratio by Brp-LPA, ROCK inhibitor Y27632 (10 μM) or myosin II inhibitor blebbistatin (10 μM) in *PLPP3* knock-downed HAEC under UF (N=4-9). **(D)** Increased *PLPP3* mRNA in intima but not in media (M) and adventitia (A) (N=3) in the PCL-artery in mice injected with nanoparticles containing *CDH5-PLPP3* plasmids. Reduced endothelial nuclear YAP by *PLPP3* overexpression. **(E)** Significantly reduced atherosclerosis and endothelial expression of CTGF and CYR61 in the PCL-artery in *Apoe*^-/-^ mice treated with nanoparticles containing *CDH5-PLPP3* plasmids compared to those with *CDH5*-only (Ctrl) plasmids (N=3-5). Data in 1A-C represent mean ± SD while mean ± SEM in Fig. 1D-E. * *P* < 0.05, ** *P* < 0.005, *** *P* < 0.0005. For two groups, unpaired student t-test was used. One-way ANOVA was used for multiple groups.

Our *in vitro* mechanistic investigations discovered an unrecognized molecular link between an atherosclerotic GWAS gene and YAP/TAZ at the interface of mechano-transduction. To establish a proof of principle to spatially manipulate *PLPP3* and YAP/TAZ signaling in vascular endothelium and treat atherosclerosis *in vivo*, we devised a method integrating nano-size carriers and endothelium-specific *PLPP3*-expressing plasmids. Nanoparticles were formulated by complexing positively-charged polyethylenimine (PEI) and negatively-charged plasmids, in which *PLPP3* was driven by a ~2.2 kb endothelium-specific *CDH5* (VE-Cadherin) promoter. Nanoparticles encapsulating *CDH5-PLPP3*-expressing plasmids, but not the control plasmids, significantly increased *PLPP3* in endothelium-enriched intima isolated from the partially carotid ligated (PCL)-artery without affecting *PLPP3* in the media/adventitia, showing endothelium-specific overexpression (Fig. 1D). Endothelial nuclear YAP in the PCL-artery was reduced by *CDH5-PLPP3*-expressing plasmids (Fig. 1D). Nanoparticle-mediated endothelial *PLPP3* restoration in the PCL-artery significantly attenuated DF-induced CTGF and CYR61 expression, and reduced atherosclerosis in *Apoe*^-/-^ mice (Fig. 1E).

Emerging results demonstrate that many GWAS genes have tissuespecific effects, underscoring the importance to integrate newly discovered GWAS mechanisms with targeted drug delivery systems. Here we demonstrate a proof of concept to spatially manipulate the atherosclerotic GWAS gene *PLPP3* in vascular endothelium where *PLPP3* uniquely responds to hemodynamics. Moreover, our results discovered a novel mechanistic link between atherosclerotic GWAS and YAP/TAZ regulation at the interface of endothelial mechano-transduction. *In vivo* restoration of endothelial *PLPP3* by targeted nanomedicine significantly inactivated YAP/TAZ signaling and reduced DF-induced atherosclerosis (Fig. 1F).

## Notes

### Competing Interest Statement

The authors have declared no competing interest.

